# Accurate estimation of intraspecific microbial gene content variation in metagenomic data with MIDAS v3 and StrainPGC

**DOI:** 10.1101/2024.04.10.588779

**Authors:** Byron J. Smith, Chunyu Zhao, Veronika Dubinkina, Xiaofan Jin, Liron Zahavi, Saar Shoer, Jacqueline Moltzau-Anderson, Eran Segal, Katherine S. Pollard

**Affiliations:** The Gladstone Institute of Data Science and Biotechnology, San Francisco, CA, USA; Chan-Zuckerberg Biohub San Francisco, San Francisco, CA, USA; Department of Biomedical Engineering, University of Calgary, Calgary, AB, Canada; Department of Computer Science and Applied Mathematics, Weizmann Institute of Science, Rehovot, Israel; Department of Molecular Cell Biology, Weizmann Institute of Science, Rehovot, Israel; Department of Gastroenterology, University of California, San Francisco, CA, USA; Benioff Center for Microbiome Medicine, Department of Medicine, University of California San Francisco, San Francisco, CA, USA; Department of Epidemiology and Biostatistics, University of California, San Francisco, CA, USA

## Abstract

Metagenomics has greatly expanded our understanding of the human gut microbiome by revealing a vast diversity of bacterial species within and across individuals. Even within a single species, different strains can have highly divergent gene content, affecting traits such as antibiotic resistance, metabolism, and virulence. Methods that harness metagenomic data to resolve strain-level differences in functional potential are crucial for understanding the causes and consequences of this intraspecific diversity. The enormous size of pangenome references, strain mixing within samples, and inconsistent sequencing depth present challenges for existing tools that analyze samples one at a time. To address this gap, we updated the MIDAS pangenome profiler, now released as version 3, and developed StrainPGC, an approach to strain-specific gene content estimation that combines strain tracking and correlations across multiple samples. We validate our integrated analysis using a complex synthetic community of strains from the human gut and find that StrainPGC outperforms existing approaches. Analyzing a large, publicly available metagenome collection from inflammatory bowel disease patients and healthy controls, we catalog the functional repertoires of thousands of strains across hundreds of species, capturing extensive diversity missing from reference databases. Finally, we apply StrainPGC to metagenomes from a clinical trial of fecal microbiota transplantation for the treatment of ulcerative colitis. We identify two *Escherichia coli* strains from two different donors that are both frequently transmitted to patients, but have notable differences in functional potential. StrainPGC and MIDAS v3 together enable precise, intraspecific pangenomic investigations using large collections of metagenomic data without microbial isolation or de novo assembly.

## Introduction

In both diseased and healthy individuals, distinct strains of the same microbial species can differ in medically relevant traits, including metabolic capacity (Joglekar et al. 2018), immunological interactions (Yang et al. 2020; Carrow et al. 2020), antimicrobial resistance (Ray et al. 2017), and pathogenic potential (Pakbin et al. 2021). Evaluating the functional potential encoded in each genome is the first step in predicting strain-specific impacts on human health, and methods that accurately determine gene content from metagenomic data can greatly improve our understanding of the extent and importance of this intraspecific diversity. Widely used tools for analyzing metagenomic data can accurately quantify the abundance of species present in a microbial community, but often fall short in characterizing variation in gene content between strains (Plaza Oñate et al. 2018). As a result, it is challenging to study the functional consequences of strain-level variation in the gut microbiome.

The most common way to study microbial gene content *in situ* is to quantify the gene families present in shotgun metagenomes, an approach referred to as “pangenome profiling”. Pangenome profiling estimates the mean sequencing depth—sometimes called vertical coverage—of a gene family as the mean number of reads aligning to each base of a representative sequence (Milanese et al. 2019). (For brevity, we use “gene” as short-hand for gene family and “depth” for mean sequencing depth throughout this paper.) Several existing tools, including PanPhlAn (Beghini et al. 2021) and MIDAS (Zhao et al. 2022a; Nayfach et al. 2016) perform pangenome profiling. However, due to several sources of error in quantifying gene depth, a second algorithm is needed to infer which genes are actually present in a specific strain’s genome, a step that we call gene content estimation. Since this strain is never directly observed in isolation—indeed, it is only a hypothesis—we refer to it as an inferred strain. Tools for gene content estimation are often based on the assumption that all encoded genes will be at a similar depth: the same as the overall species depth (Plaza Oñate et al. 2018), which can be directly estimated from the depth of species marker genes (Blanco-Míguez et al. 2023; Milanese et al. 2019). Therefore, the depth ratio—the ratio of a given gene’s depth to the overall species depth in a sample—can be used as the key criterion for the assignment of genes to a species (Nayfach et al. 2016).

However, gene content estimation using pangenome profiles faces four key challenges:

1. an incomplete set of representative gene sequences in pangenome reference databases,
2. ambiguous alignment of short-reads to multiple sequences both within and across species (“cross-mapping”),
3. poor discrimination between present and absent genes for species at low depth due to high variance of the depth ratio, and
4. a decreased depth ratio for strain-specific genes when other strains of the same species are also abundant (“strain mixing”).

Significant progress towards (1) has been recently achieved by expanding pangenome reference databases to include metagenome assembled genomes (MAGs), substantially improving their coverage for human gut species (Almeida et al. 2020). Unfortunately, MAGs can contain cross-species contamination and genome assembly errors such as gene fragmentation. The frequency and types of these errors varies depending on the source and quality of the MAGs, as well as whether they were assembled from short-read or long-read sequencing data. Both types of errors can exacerbate cross-mapping (2), potentially reducing the accuracy of pangenome profiling (Zhao et al. 2023). Careful curation of the pangenome database is needed to reduce the impact of these issues. One promising approach for dealing with low depth (3), is to combine data across multiple samples (Carr et al. 2013; Plaza Oñate et al. 2018), taking advantage of increased depth from pooling reads. As a bonus, the correlation between the species depth and gene depth can be used as an additional criterion to better exclude genes with cross-mapping (2). However, combining samples can exacerbate the impacts of strain mixing (4). Methods are needed for strain-aware gene content estimation that benefit from the increased sensitivity and specificity of multiple samples while also accounting for intraspecific variation.

Here we introduce StrainPGC (“Strain Pure Gene Content”), a computational method designed to accurately estimate the gene content of individual microbial strains. StrainPGC leverages modern strain tracking tools to separate samples into strain-pure subsets in order to combine data from pangenome profiling across multiple metagenomes. We also describe changes in MIDAS v3, including updates to the pangenome database (MIDASDB) and profiling pipeline to reduce cross-mapping, improve quantification, and facilitate the interpretation of strain-specific gene content. As part of a complete workflow, our method requires only shotgun metagenomes as input and outputs estimates of the gene content of individual strains. After validating our workflow with a complex synthetic community (hCom2) (Jin et al. 2023), we use it to explore strains in the Human Microbiome Project 2 (HMP2) (Proctor et al. 2019) and in a trial of fecal microbiota transplantation for ulcerative colitis (UCFMT) (Smith et al. 2022b). We find novel strain diversity not captured in existing reference databases as well as widespread variation in gene content, including functions with likely clinical relevance.

## Results

### Updated pangenome profiling and strain-specific gene content estimation

MIDAS v3 represents a major upgrade to the pangenome profiling pipeline intended to improve the completeness, curation, and interpretability of gene abundance estimates. We updated the pangenome database construction and gene annotation process as well as the profiling algorithm, and describe these in the Methods section below. In each sample, MIDAS quantifies the depth of the genes in each species’ pangenome.

In order to further refine these pangenome profiles for individual samples into gene content estimates for specific strains, we designed StrainPGC, a novel method that integrates data from multiple metagenomes to overcome the limitations of pangenome profiling for characterizing intraspecific variation (Fig. 1A-C). For each species, StrainPGC takes in pangenome profiles and two other inputs, a list of species marker genes, and a list of “strain-pure” samples for each of the desired strains. The StrainPGC algorithm can be summarized as follow: First, the species depth in each sample is estimated based on mean depth of the provided marker genes. Next, based on this depth, “species-free” samples are identified as those where the species is below a minimum detection limit (in this work 0.0001x). Then, separately for each strain, two statistics are calculated for each gene (Fig. 1C): (1) the depth ratio is the total gene depth divided by the total species depth across that strain’s pure samples; (2) the correlation score is the Pearson coefficient between the gene’s depth and the overall species depth across both this strain’s pure samples and the species-free samples. For each strain, genes passing a minimum threshold for both of these statistics—the depth ratio and the correlation score (Fig. 1D)—are estimated to be present in that strain’s genome. Finally, two quality control statistics (described below) are calculated for each strain intended to flag those likely to be of low accuracy.

**Figure 1:**
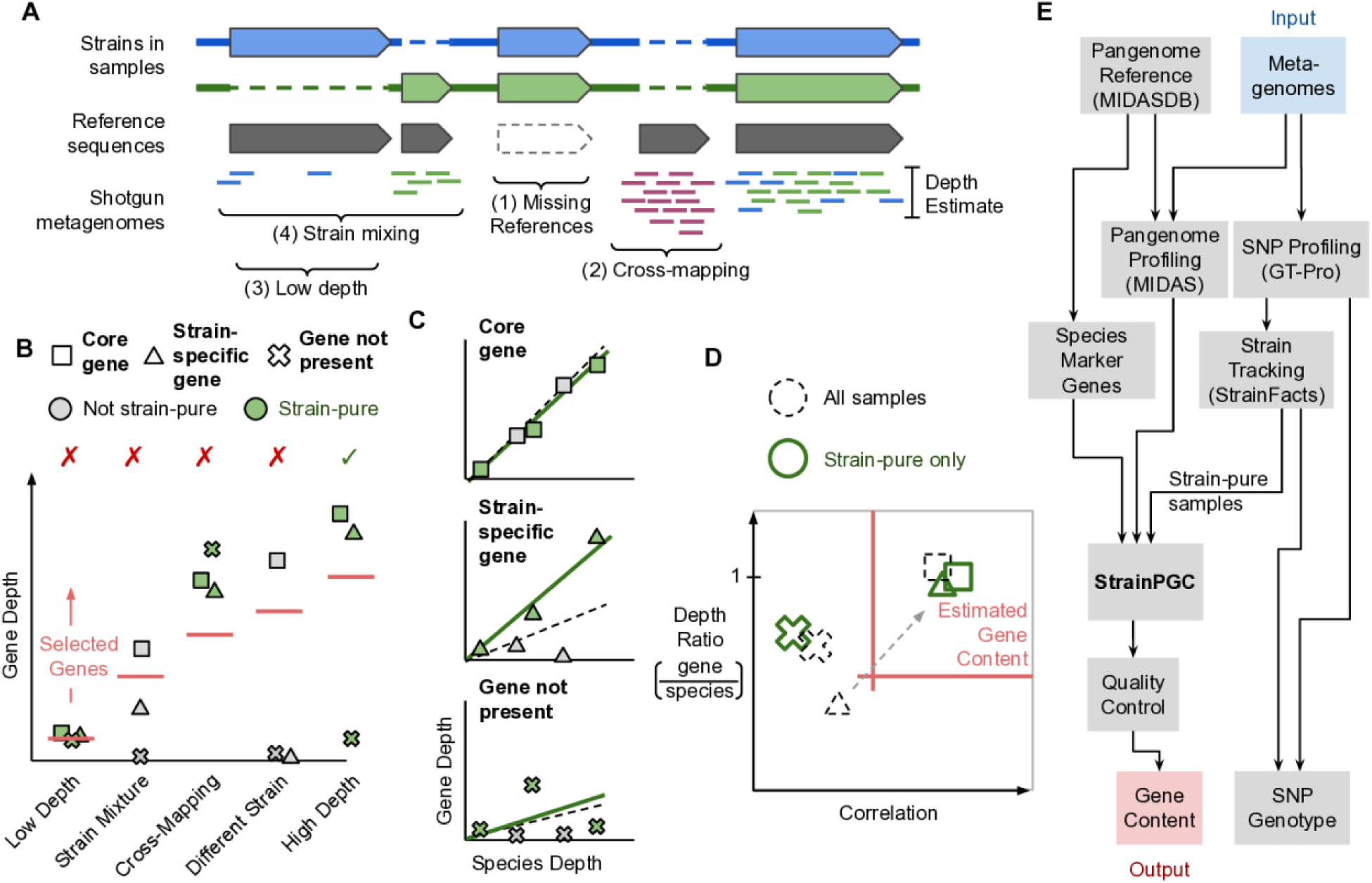
Conceptual overview of strain-resolved gene content reconstruction using StrainPGC. **(A)** Schematic representation of pangenome profiling, which estimates gene depth based on short-read alignment. The illustration represents profiling of a hypothetical microbial population harboring two strains of the same species (blue and green), each with both shared and strain-specific gene content. Four key challenges for pangenome profiling and gene content estimation are highlighted (brackets, see Introduction). **(B)** Limitations of gene content estimation using single samples. Depth is shown across five samples for three genes: one gene is ubiquitous across strains (“core”), another is found in only the strain of interest (“strain-specific”), and a third not present in the strain of interest but is susceptible to cross-mapping (“not present”). Samples are separated along the x-axis and represent five characteristic scenarios: a sample where the species is not deeply sequenced (“low depth”), a sample with multiple strains of the species (“strain mixture”), a sample exhibiting erroneous depth due to read mapping errors (“cross-mapping”), a sample with an entirely different strain of the species (“different strain”), and a high depth, strain-pure sample (“high depth”). Colors distinguish between strain-pure samples (green markers) and samples with a different strain or a mixture of more than one strain (gray markers). Traditional, single-sample analysis estimates gene content by selecting genes with a minimum depth (red, horizontal line, which is chosen based on the species’s depth). As a result, samples with low depth, cross-mapping, and strain mixing all lead to decreased accuracy (indicated with red x’s). Only gene content estimation in a strain-pure, high-depth sample without cross-mapping (green check) accurately reflects the strain of interest. **(C)** Relationship between gene depth and species depth for each of the three genes (panels) across the five samples (marker shape and color as in B). For each, the linear relationship is shown between species depth and gene depth in the set of strain-pure samples (solid green line). We contrast this fit with the linear relationship across all five samples without considering strain variation (dashed line). **(D)** Schematic depiction of how StrainPGC estimates gene content based on both correlation and depth ratio. The red lines indicate the thresholds of depth ratio and correlation used by StrainPGC to select genes. With all samples combined (dashed markers), the “not-present” gene is correctly excluded due to low correlation, and the core gene is correctly included, but the strain-specific gene is lost due to its low depth ratio and correlation. Analyzing the strain-pure set separately moves the strain-specific gene into the selection region (dashed arrow), increasing accuracy. **(E)** Schematic depiction of our integrated workflow to infer gene content across strains using only shotgun metagenomic reads as input.

While StrainPGC is designed to accept strain-pure samples identified using a variety of strain tracking approaches, in this work we apply GT-Pro (Shi et al. 2022), an assembly-free algorithm for tallying single-nucleotide polymorphisms (SNPs) in shotgun metagenomic reads, followed by StrainFacts (Smith et al. 2022a), which harnesses these SNP profiles to precisely identify individual strains within species and quantify their relative abundances. For each species, we consider samples estimated to be ≥ 95% the majority strain as pure. Strains analyzed in this work are therefore defined based on their SNP-genotypes, with gene content estimated as a subsequent step.

StrainPGC is open source and freely available at https://github.com/bsmith89/StrainPGC. We integrated pangenome profiling, strain tracking, and gene content estimation into a complete Snakemake (Mölder et al. 2021) workflow (Fig. 1E) intended for studying the human gut microbiome. As with other tools, the computational resources required to run the full pipeline may be substantial and are dominated by the requirements for read alignment with Bowtie2 (Langmead and Salzberg 2012). By comparison, even for large datasets, the StrainPGC core algorithm generates results for all strains of a species and requires only a few GBs of RAM at peak (see GitHub README for details). Our analysis harnesses the comprehensive Unified Human Gastrointestinal Genome (UHGG) reference collection (Almeida et al. 2020) and requires only raw metagenomic reads as user-provided input.

### StrainPGC accurately estimates gene content of strains in a complex synthetic community

In order to evaluate StrainPGC’s performance, we ran our workflow on 276 publicly available metagenomes derived from experimental manipulations of the hCom2 synthetic bacterial community (Jin et al. 2023). The shared inoculum was composed of 117 bacterial isolates spanning 8 phyla, each with a high-quality genome assembly, which we refer to as ground truth genomes (Fig. 2A). Most species were represented by a single strain, some by 2 or 3 strains, and one by 4 (Fig. 2B). We refer to the collection of ground-truth genomes and experimental metagenomes as the hCom2 benchmark dataset. We annotated predicted protein-coding genes in the ground truth genomes with EggNOG OGs (Fig. 2A). After removing species that could not be genotyped by GT-Pro, or that were undetected in metagenomes, the benchmarking task amounted to 87 species encompassing 97 strains and with highly disparate depths (estimated maximum sample depth interquartile range of 2.7–22.4x) (Fig. 2B). We applied StrainPGC to estimate gene content across inferred strains, matched each ground truth strain to a single inferred strain based on SNP genotypes, and compared the EggNOG OGs annotations between these. In this benchmark, StrainPGC had a median precision of 0.96 (IQR: 0.90–0.98; Fig. 1C), a recall of 0.88 (0.82–0.93), and an F1 score of 0.91 (0.87–0.94).

**Figure 2:**
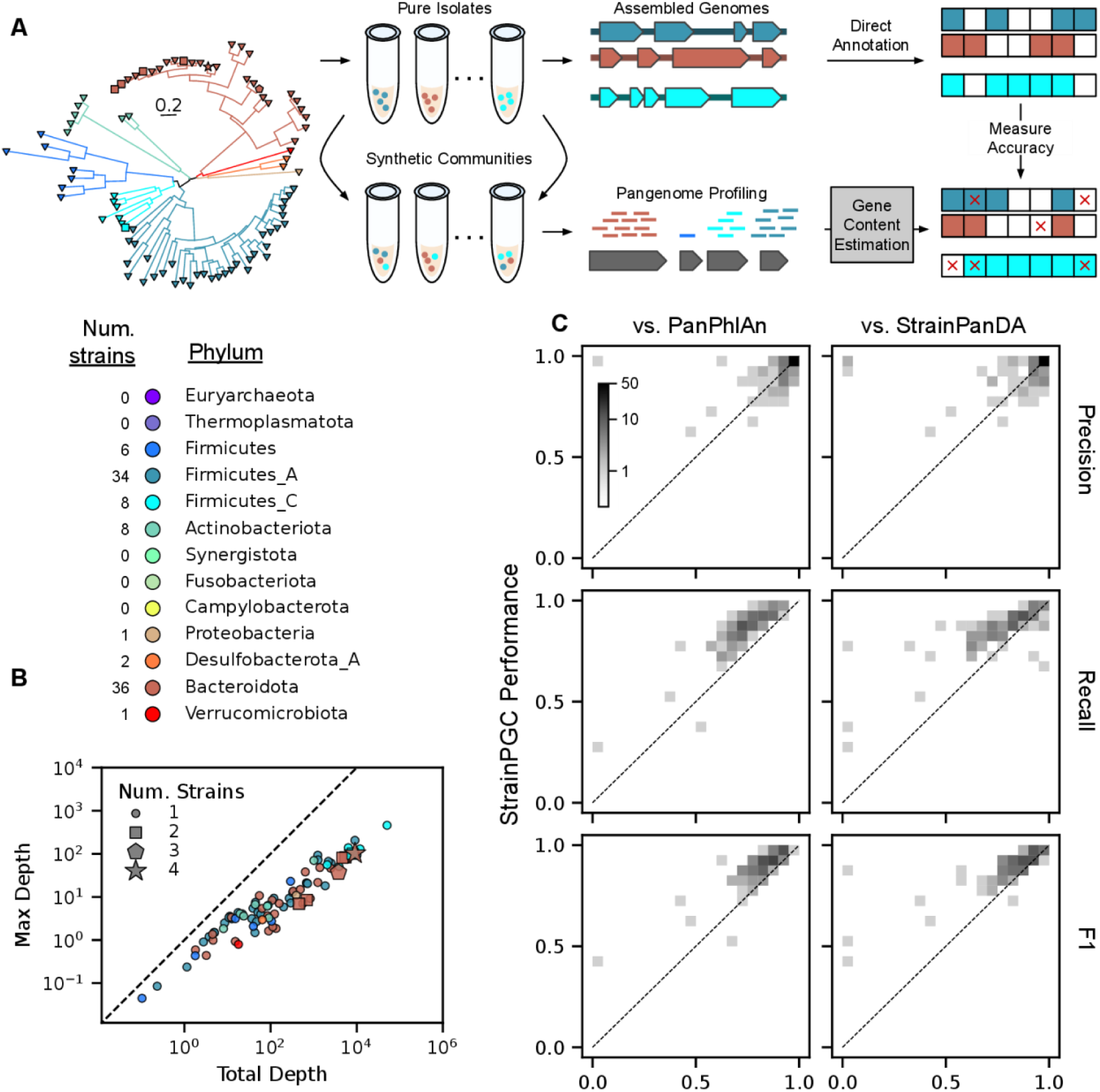
Evaluation of StrainPGC’s gene content estimation performance on a highly diverse, synthetic community (Jin et al. 2023). **(A)** Schematic diagram of our procedure for benchmarking gene content estimates using the hCom2 synthetic community constructed to reflect the species and strain diversity found in human gut microbiomes (Cheng et al. 2022). StrainPGC and alternative tools were applied to pangenome profiles from different samples derived from the synthetic community, and estimates of gene content were compared to high-quality reference genomes for 97 strains. Strains were drawn from 95 species across 8 phyla (phylogenetic tree on the left, colored by phylum, scale bar in units of substitutions per position). **(B)** Core genome depths of 87 detectable benchmarking species span more than two orders of magnitude. Points represent individual species, are colored by phylum, and are placed based on that species’s maximum depth across samples (x-axis) and total depth summed over all samples combined (y-axis). Species are closer to the 1-to-1 diagonal (dashed line) when the sample with the highest depth contributes more of their total depth. Some species are represented by more than one strain (marker shape). **(C)** Accuracy of gene content estimates by StrainPGC (y-axis) compared to PanPhlAn (Beghini et al. 2021) and StrainPanDA (Hu et al. 2022) (x-axes), as measured by precision, recall, and F1. All three indices range between 0 and 1, and higher values reflect better performance. The data are represented as two-dimensional histograms using a gray density scale to represent the number of strains falling in each (x, y) bin; density above the 1-to-1 diagonal (dotted line) indicates strains where StrainPGC outperformed the alternative on that index. The relationship between performance and strain sequencing depth or sample number are shown in Supplementary Figure S1.

We next compared StrainPGC’s performance to two alternative, state-of-the-art methods: PanPhlAn (Beghini et al. 2021), which is widely used and operates on single samples, and StrainPanDA (Hu et al. 2022), a recently published tool that harnesses information across multiple samples and applies non-negative matrix factorization to jointly estimate gene content and strain depth (Fig. 2C). For all three methods, we used the same reference database (UHGG) and pangenome profiles as input, thereby comparing the gene content estimation approaches on an equal basis. However, since strains inferred using PanPhlAn and StrainPanDA do not have SNP genotypes to be used for matching, for each hCom2 genome, we instead selected the inferred strain with the highest F1 score, giving these two methods an advantage. Nonetheless, StrainPGC performed better on average than either alternative: a median increase of 0.069 in F1 score compared to PanPhlAn (IQR: 0.038–0.093; p < 1e-10 by Wilcoxon, non-parametric, paired, t-test) and 0.042 relative to StrainPanDA (IQR: 0.022–0.079; p < 1e-10). All three tools had similarly high precision, and the superior performance of StrainPGC was driven primarily by a dramatic reduction in the false negative rate (FPR: 1 -recall): a median of just 49% of PanPhlAn’s and 60% of StrainPanDA’s FPR.

For all three tools, strains with higher estimated depth had better performance on this benchmark (Spearman’s correlation between maximum strain depth across samples and F1 score: Spearman’s ⍴ = 0.29, 0.55, and 0.32 for StrainPGC, PanPhlAn, and StrainPanDA, respectively; Supplementary Figure S1). We also find a correlation between the number of strain-pure samples and F1 for all three tools (⍴ = 0.33, 0.42, and 0.34, respectively, Supplementary Figure S1). Interestingly, StrainPGC’s precision was less tightly related to depth than either PanPhlAn or StrainPanDA (⍴ = 0.19, 0.54, and 0.55, respectively). Since we controlled for the upstream pangenome profiling, these findings support the use of the Pearson correlation across strain-pure samples as a filtering criterion for gene content estimation, allowing StrainPGC to maintain high precision even while greatly increasing recall. In particular, we find our approach upholds this specificity—even at low depths—more effectively than existing methods and that performance was fairly stable for strains with ≥ 5 samples, or when at least one sample had depth ≥ 1x (Supplementary Figures S1).

In real-world applications—where ground-truth gene content is not known a priori—it is beneficial to understand the confidence of StrainPGC estimates. We, therefore, calculated two scores to serve as proxies for accuracy and compared these to the performance we measured on the hCom2 datasets. First, we hypothesize that the fraction of high-prevalence, species marker genes assigned to a given inferred strain reflects the overall completeness of the estimated gene content for that strain. Indeed, across strains in the hCom2 benchmark, we found a strong correlation between the fraction of species marker genes and the F1 score (⍴ = 0.60, p < 1e-10). As expected, this appears to be driven primarily by a strong association with the recall (⍴ = 0.63, p < 1e-10); a weaker correlation was found with the precision (⍴ = 0.34, p < 1e-3). Second, for strains suffering from low signal-to-noise, such as those at low sequencing depths, the depth ratio of assigned genes will be more variable. We, therefore, calculated a noise index reflecting: the standard deviation across all assigned genes of the log10-transformed depth ratio. For this score, we found a negative correlation with the F1 score (⍴ = - 0.68, p < 1e-10), this time driven by an association with the precision (⍴ = -0.58, p < 1e-9) as well as recall (⍴ = -0.53, p < 1e-8). In our benchmark, the 22 strains with < 95% species marker genes or a noise index > 0.25 had substantially lower F1 scores than those that passed this quality control (median of 0.83 versus 0.92, p < 1e-5 by MWU test). We propose using these two criteria together in order to exclude inferred strains with lower accuracy gene content estimates.

### Inferred strains in publicly available metagenomes substantially expand the catalog of intraspecific diversity

We applied our workflow to the 106 subjects and 1338 samples of the HMP2 metagenome collection—which we refer to as simply the HMP2 throughout this paper.

First, to explore the strain-level diversity that might be discovered in publicly available datasets, we used StrainFacts to identify and estimate the distribution of strains based on SNP profiles. We defined detection as an estimated depth of ≥ 0.1x, a threshold chosen to balance false positives with the sensitivity of strain tracking. All species combined, a median of 59 strains were detected in each metagenomic sample and 191.5 across all samples from each subject (Fig. 3A). This strain-level diversity was highly subject-specific; among inferred strains detected in two or more samples, 36% were detected in just one subject, and only 34% were detected in three or more (Fig. 3B). Strain sharing was dramatically more common in pairs of samples from the same subject than in pairs of samples from different subjects (mean of 36.7 shared, detected strains from same subject vs. 0.7 from different subjects, p < 1e-10 by MWU; Fig. 3C), consistent with prior studies of the HMP2 and other cohorts (Lloyd-Price et al. 2017).

**Figure 3:**
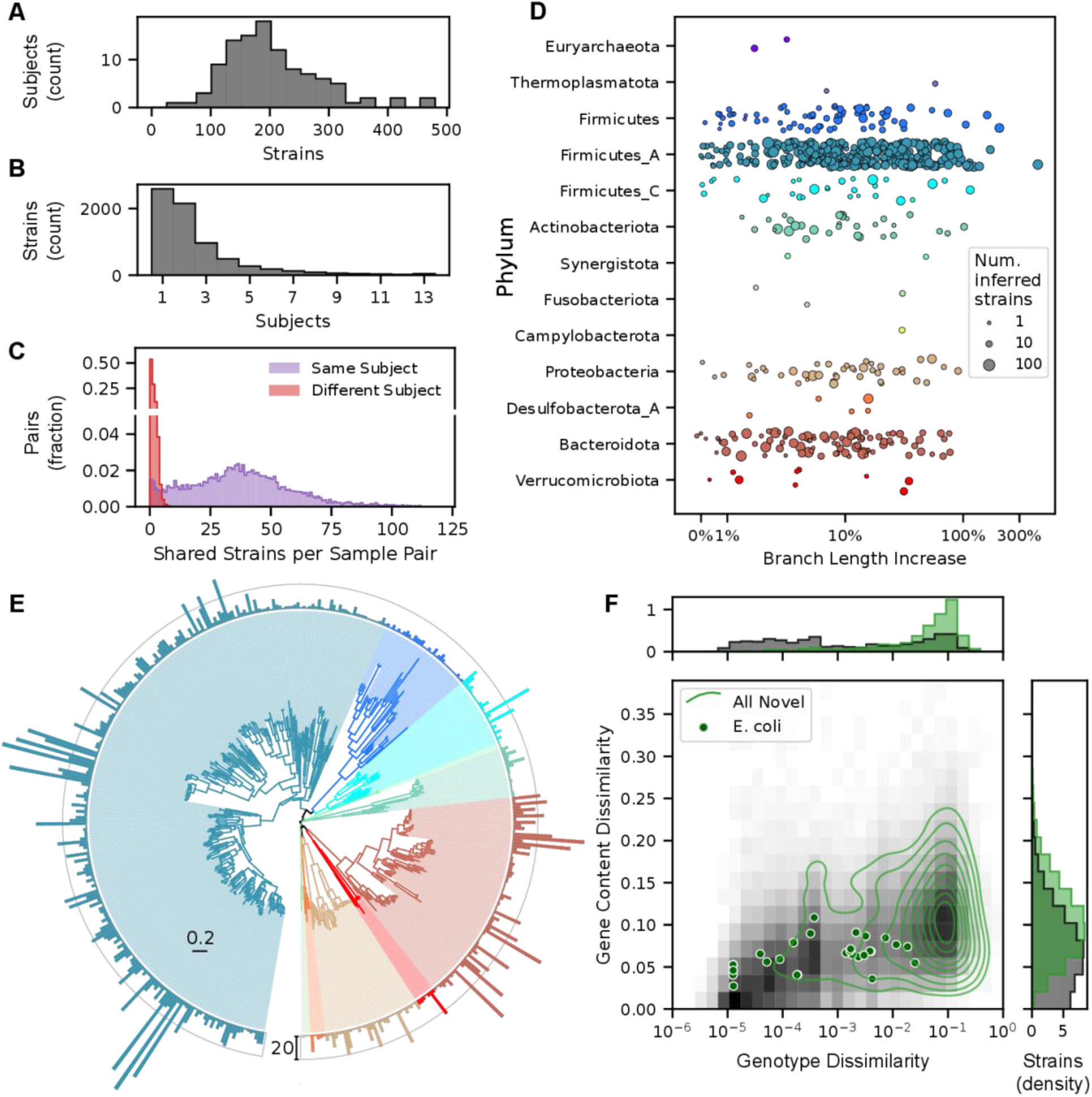
Strain diversity in the HMP2 metagenome collection. **(A–B)** BHistograms reflecting the distribution of inferred strains of any species across subjects in the HMP2 metagenome collection. **(A)** Number of strains for 106 subjects, summed over all samples (median of 11 samples per subject; IQR: 9–14). Most subjects harbor between 100 and 300 inferred strains (median of 191.5). **(B)** Number of subjects where each strain was detected. Only strains found in two or more samples are tallied. Most strains (67%) were found in just one or two subjects. **(C)** Number of strains shared in any pair of samples from the same (purple) or different (red) subjects. Pairs of samples from different subjects shared a mean of just 0.7 strains. **(D)** A substantial increase in strain diversity was captured when including inferred strains. Diversity was quantified based on total branch length in a hierarchical clustering (UPGMA) of all SNP genotypes, and the increase was measured as the change in branch length relative to a tree with only reference strains. Points represent individual species, are colored by phylum, and increasing size reflects a larger number of inferred strains. Five species with fewer than 3 inferred strains had a small decrease in branch length when inferred strains were included; one of these is excluded from the plot, left of the x-axis limit. **(E)** Taxonomic diversity of 3504 inferred strains of Bacteria. The species tree is colored by phylum as in (D). Species that had no strains with estimated gene content were omitted, and bars around the outer ring indicate the number of inferred strains (outer ring indicates 20 strains). The branch length scale bar (interior) is in units of substitutions per position. **(F)** Estimated genotype and gene content dissimilarity from the closest reference genome. Joint (main panel) and marginal distributions (panels above and to the right) are plotted for all high-quality reference (gray background) and inferred (green contours) strains of all species. Gene content dissimilarity of inferred strains is calculated after batch correction (see Methods). Points reflecting each of 28 inferred E. coli strains are also shown. Green contours in the main panel reflect deciles in the 2D kernel density estimator.

Concordant with this level of strain diversity, estimated genotypes for inferred strains were often distinct from the closest reference strain. Using SNP profiles in strain-pure samples, we estimated each inferred strain’s genotypes as the consensus allele, masking ambiguous positions. Among inferred strains with ≥ 100 genotyped positions, 68% had a genotype dissimilarity of greater than 0.05 to the closest reference. Representing the strain diversity of each species with a UPGMA tree, we calculated the increase in total branch length when including inferred strains relative to only references (Fig. 3D). For many species, a substantial increase in total branch length was observed: more than 10% for 288 species, more than 20% for 183 species, and more than 50% for 63 species when inferred strains were included. Overall, these findings suggest that inferring strains from publicly available metagenome collections will reveal novel intraspecific diversity not already found in reference databases.

To further evaluate the expected performance of StrainPGC in real-world scenarios, we performed an *in silico* experiment, using five *E. coli* genomes not in the UHGG reference collection (Davidova-Gerzova et al. 2023), spiking-in simulated reads to the HMP2 dataset. These benchmark genomes represent a range of divergence from the closest reference genome similar to what we found for the inferred strains. Despite this additional complexity and reference bias, we observed F1 scores equivalent to those in the hCom2 benchmark (Supplementary Material and Supplementary Table S2).

Having in this way validated its performance in the HMP2 dataset, we next applied StrainPGC to the novel, inferred strains. After quality control, we estimated gene content for 3511 strains in 443 species across 12 phyla (Fig. 3E). Strains had a median of 9 strain-pure samples (IQR: 5 - 13). While these were primarily Bacteria, we were also able to estimate gene content for strains in three species of Archaea. The largest number of inferred strains were classified in the phylum Firmicutes_A (2232 strains; an additional 80 and 141 strains were also in “Firmicutes”, and “Firmicutes_B”, respectively, which are classified as separate phyla in the GTDB taxonomy), followed by Bacteroidota (727), and Proteobacteria (189). Hence, StrainPGC resolved gene content for myriad strains across a diverse set of species found in the human gut (Supplementary Table S1).

Just like SNP genotypes, for most inferred strains, the estimated gene content was quite distinct from the closest reference. Measuring dissimilarity using the cosine dissimilarity after batch correction (see Methods), inferred strains were a median of 0.18 from the closest, high-quality reference genome (Fig. 3F). As would be expected, strains with more dissimilar SNP genotypes were often those with dissimilar gene content as well. For instance, across the 28 inferred strains of *E. coli*, we found a significant correlation between the gene content dissimilarity and the genotype dissimilarity (Spearman’s ⍴ = 0.44, p = 0.018; Fig. 3F). This suggests that the increased diversity captured by StrainPGC facilitates expanded analyses of intraspecific gene content variation in the gut microbiome.

### Estimated gene content enables pangenome analyses in prevalent human gut microbes

To demonstrate the value of gene content estimates derived from the HMP2 for pangenome analysis, we focused on the 99 species with estimated gene content for 10 or more inferred strains (Median: 17 inferred strains per species, IQR: 12–28, 7 phyla). For each species, we calculated the prevalence and distribution of genes across strains. Gene prevalence estimates based on inferred strains were highly correlated with the prevalence observed in high-quality reference genomes (r = 0.84, p < 1e-10; Fig. 4A), supporting the consistency of our estimates with the existing reference database.

**Figure 4:**
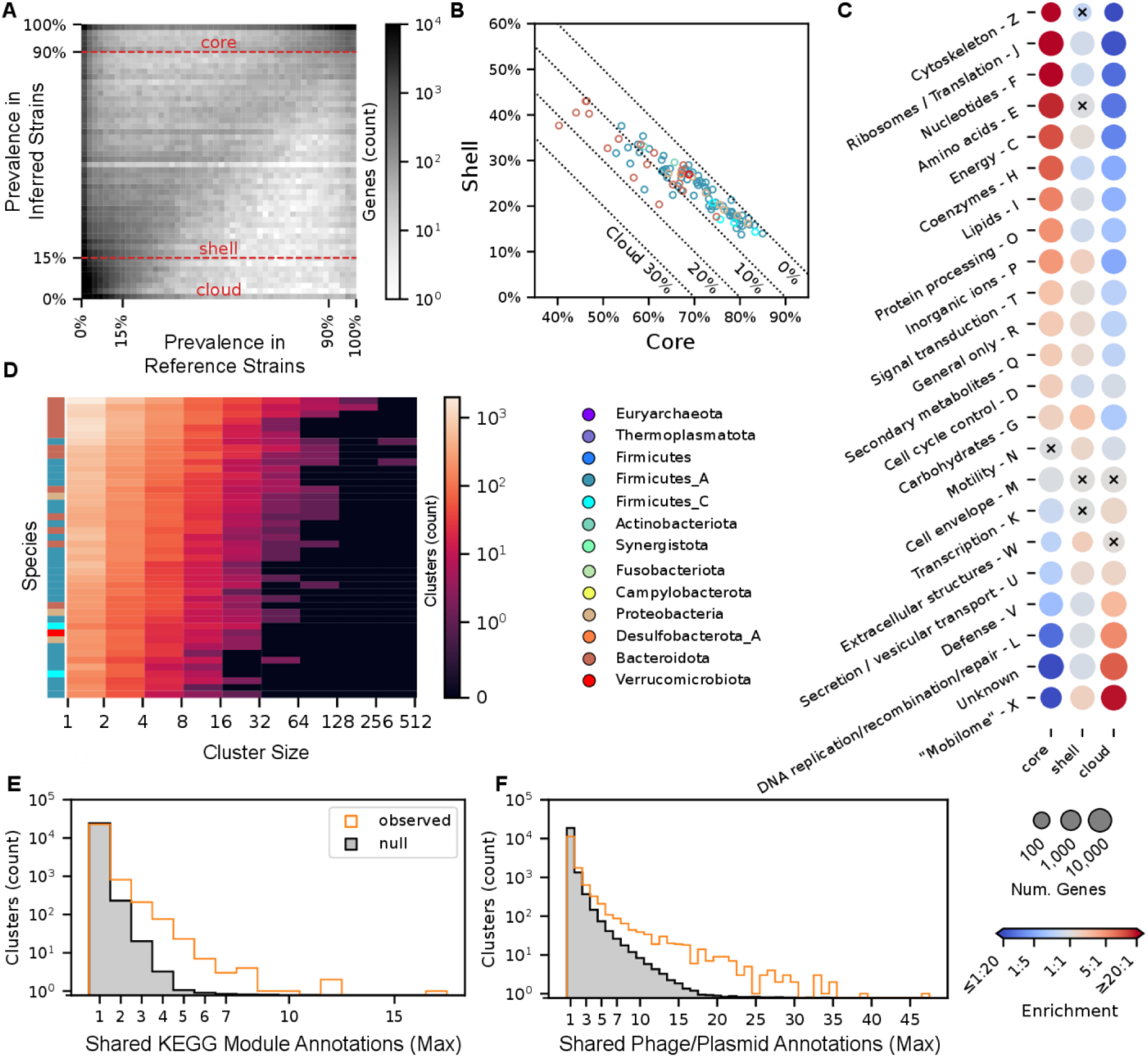
StrainPGC reveals patterns of gene content variation across dozens of species. **(A)** Gene prevalence across inferred strains from HMP2 is very similar to prevalence in reference genomes. Combining genes from all species, the 2D histogram shows the joint distribution of prevalence estimated from reference genomes (x-axis) and inferred strains (y-axis). These independent estimates are highly concordant, with higher density along the diagonal. Dashed horizontal lines represent the thresholds defining core, shell, and cloud prevalence classes based on inferred strains. **(B)** Fraction of shell versus core genes in inferred strains. For each species (circle), x and y values are the median gene content in the core and shell classes, respectively. The remaining gene content is composed of cloud genes and is indicated by the dotted diagonal lines. Markers are colored by phylum. Analogous results calculated using reference genomes are shown in Supplementary Figure S2. **(C)** Enrichment (red) or depletion (blue) in genes of various functional categories in each of the core, shell, and cloud prevalence classes. Dots representing each COG category (rows) and prevalence class (columns) are colored by odds ratio, with red and blue indicating enrichment and depletion, respectively. Dot size reflects the number of genes in that prevalence class that are in the given functional category. All enrichments/depletions shown are significant (Two-tailed Fisher Exact Test; p < 0.05), except for those marked with a black cross. COG categories A, B, and Y are omitted, as these had very few members (173, 74, and 0 genes, respectively). **(D)** Gene co-occurrence clusters based on estimated gene content. The heatmap depicts histograms for each of 44 species (rows) of cluster sizes (columns). Colors indicate the number of clusters in each interval, and labels along the x-axis indicate the bounds of the intervals (left exclusive, right inclusive). Colors on the left indicate phylum as elsewhere. **(E, F)** The maximum number of related annotations in each co-occurrence cluster. The orange histogram represents the observed distribution, while the gray region is the mean in each bin across 100 random permutations of cluster labels (i.e. the null distribution). The higher number of clusters with multiple, shared annotations in the observed data compared to the null suggests clumping of **(E)** KEGG module and **(F)** phage or plasmid genes into co-occurrence clusters.

Based on these de novo prevalence estimates, we assigned genes to the “core” (≥ 90% prevalence), “shell” (< 90% and ≥ 15%), or “cloud” (< 15%) pangenome fractions. We then calculated the portion of estimated gene content that fell into each prevalence class for each inferred strain (Fig. 4B). Computing the median first within and then across species, genes in the core fraction made up 70% (IQR: 63–76%) of each strain’s estimated gene content, shell fraction 25% (19–28%), and cloud fraction 5% (4–9%), in general agreement with reference genomes (Supplementary Figure S2). Certain categories of functional annotations were more common in each fraction (Fig. 4C). Core genes were enriched for COG categories with housekeeping functions while the cloud pangenome was enriched in functional categories including the mobilome, and defense mechanisms. The COG category for DNA replication, recombination, and repair and genes without a COG category were also enriched in the cloud pangenome, possibly indicating that many of these genes are also related to the mobilome. Broadly, these patterns of enrichment confirm our expectations that core genes perform obligate functions and make up a plurality of genes for most strains.

We also identified 200 genes that were annotated with antimicrobial resistance functions. Of these, 168 were in the cloud, 32 were in the shell, and none were in the core pangenome fraction. Across all 3511 high-quality strains, 482 (14%) of these had at least one gene with an AMR annotation. We also found differences across phyla in the fraction of strains with at least one annotation. For Bacteroidota, 37% of strains (271 of 727) had an AMR gene, as did 22% of Proteobacteria (41 of 189). However, only 9% of Firmicutes (7 of 80), 7% of Firmicutes_A (151 of 2232), 6% of Actinobacteria (4 of 64) and 4% of Firmicutes_C (6 of 141) had an annotation. These results are consistent with our expectation that resistance mechanisms are highly variable within species and more common in gram-negative bacteria.

As an assembly-free approach, gene content estimation lacks synteny information, which can be useful for understanding biological phenomena such as operonic co-regulation and horizontal gene transfer. To get around this limitation, we clustered genes based on the Pearson correlation of their presence and absence across inferred strains in the HMP2. For the 44 species with more than 20 high-quality inferred strains, we identified 36,208 co-occurring gene clusters with 2 or more members, a median of 681.5 per species (Fig. 4D). Genes in the same cluster were more likely to have related annotations; clusters having three or more genes in the same KEGG module were 12.7x more common than expected by random chance (n = 100 permutations of cluster labels within species, p < 1e-2; Fig. 4E). Likewise, phage-or plasmid-associated genes were more frequently found in the same clusters than expected by chance (three or more shared annotations 2.4x more common, p < 1e-2; Fig. 4F). This supports our interpretation of StrainPGC–enabled gene co-occurrence clustering across genomes as evidence of related biochemical function or linked transmission, which may help to generate testable hypotheses about relationships between genes in a species’ pangenome. Overall, large surveys of gene content estimated by StrainPGC have the potential to vastly expand the coverage and diversity of pangenome analyses.

### Integrative analysis of *E. coli* strain gene content can inform the selection of donors for fecal microbiota transplantation

We next sought to assess the potential utility of StrainPGC gene content estimates for optimizing microbial therapies such as FMT. Current donor screening protocols focus on detection of known pathogens and do little to match donors to recipients or optimize for transmission and engraftment of particular microbial functions. To assess the sensitivity of our approach for comparing donor strains, we re-analyzed metagenomes from a previously published study of FMT for the treatment of ulcerative colitis (Smith et al. 2022b). We refer to these metagenomes as the UCFMT dataset. As a proof-of-concept, we focused on strains of *E. coli*, a well-studied and highly prevalent member of the human gut microbiome with well-documented examples of not only pathogenic but also commensal and even probiotic strains (Blount 2015).

Using 231 samples collected longitudinally from patients (189 samples) and donors (42 samples from three of four donors) in the UCFMT study, we identified and tracked strains using StrainFacts. Focusing on D44 and D97—the two donors with the most metagenomic samples and who contributed materials to the most recipients—we observed robust, repeated transmission of strains during FMT (Fig. 5A). Next, with StrainPGC, we obtained gene content estimates for inferred strains of *E. coli*; 18 passed quality control. In order to examine their genetic relatedness—and to put them in the context of the earlier pangenome analysis—we combined inferred strains from the UCFMT and HMP2 metagenomes and generated a UPGMA tree based on their SNP genotype dissimilarity (Fig. 5B). As before, genotype and gene content were related (Fig. 5B): for the combined set of inferred strains, we found a robust correlation between the cosine dissimilarity of the shell pangenome fraction—defined above using the HMP2 strains—and genotype dissimilarity (r = 0.88, Fig. 5C).

**Figure 5:**
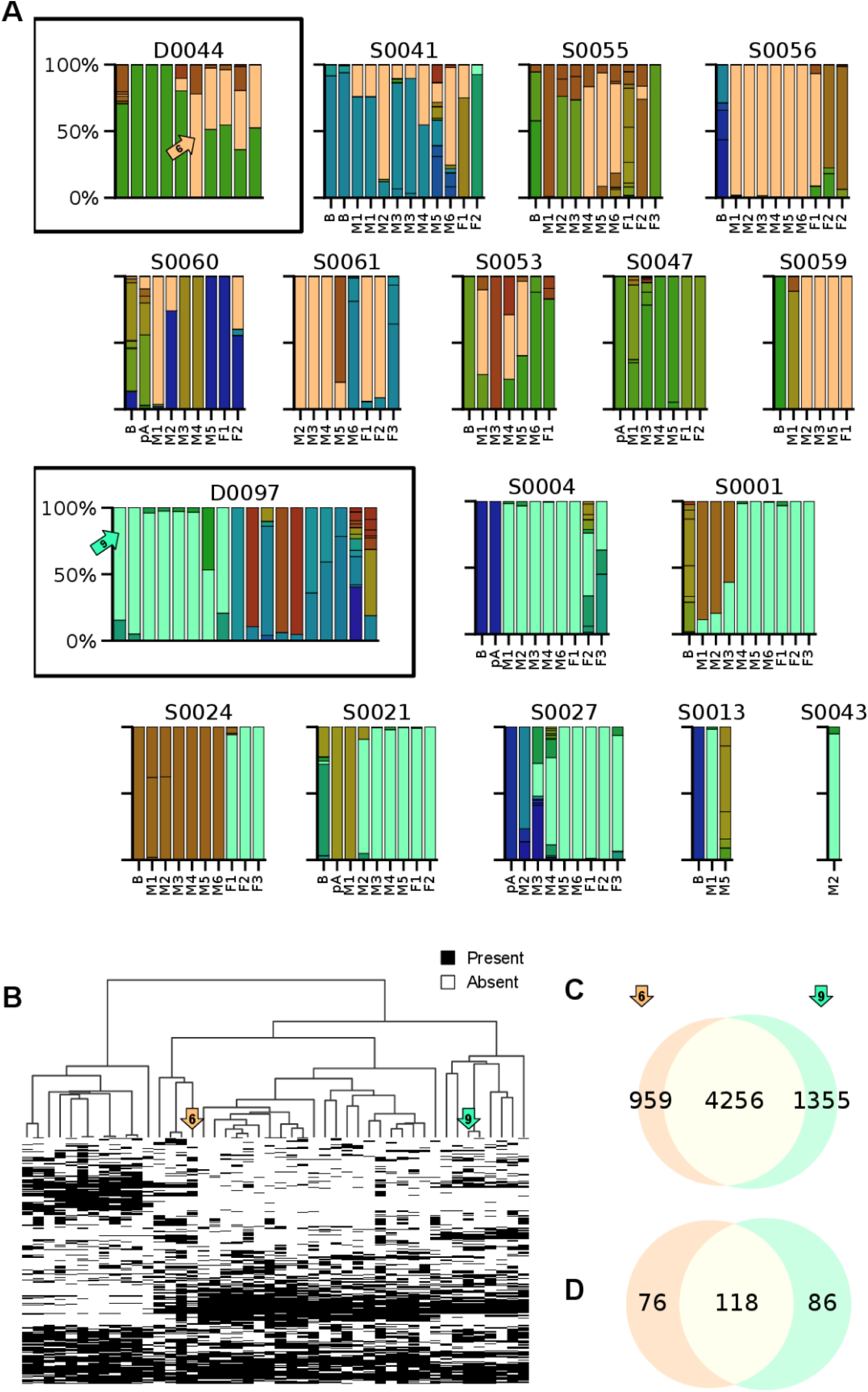
Different donors in a fecal microbiota transplant (FMT) trial (Smith et al. 2022b) have engrafting E. coli strains that differ in their functional potential. **(A)** E. coli strains found in repeated sampling of two independent donors’ fecal materials (boxed panels) and in the fecal time series of their respective recipients. Columns in each panel represent individual samples, colors represent E. coli strains inferred from StrainFacts, and the height of colored bars indicates strain abundance normalized to total E. coli abundance in the sample. For donors, samples are ordered arbitrarily. Recipient samples are ordered by collection day and include samples at baseline (labeled “B”) collected before initial FMT treatment, samples collected before each of up to six maintenance FMT doses (labeled “M1” to “M6”), and up to three follow-up samples (labeled “F1” to “F3”). For a subset of recipients, samples were also collected after antibiotic treatment and before FMT (labeled “pA”, post-antibiotics). For each donor, one strain (tan in D44, aqua in D97) showed a high rate of engraftment in recipients at follow-up. **(B)** Comparison of shell gene content between inferred strains from the FMT experiment (18 strains) and E. coli strains from the HMP2 (28). Heatmap indicates the presence and absence of genes (rows) across inferred strains (columns). Strains are ordered by UPGMA tree of estimated SNP genotype dissimilarity. Genes are filtered to only the 3,134 genes in the shell pangenome fraction. Arrows (tan and aqua) highlight the high-engraftment strains from panel (A). **(C, D)** Estimated gene content that is shared and distinct between the two high-engraftment strains. Venn diagrams depict the intersection of **(C)** genes and **(D)** gene co-occurrence clusters.

In recipients of donor D44, one strain, strain-6, stood out as frequently present both during six weeks of maintenance dosing and in subsequent follow-up sampling (Fig. 5A). Likewise, strain-9 engrafted frequently for recipients of D97. Both strains were very closely related to isolate genomes represented in the UHGG reference collection: strain-6 matched GUT_GENOME288864 (GenBank accession GCA_009896305.1) with an identical genotype at all 74,229 shared SNP positions, and strain-9 matched GUT_GENOME140932 (GCA_000408385.1) with just 7 mismatches across 79,260 shared SNP positions. The two inferred strains had a SNP genotype dissimilarity to each other of 0.23, similar to the median dissimilarity across all pairs of UCFMT strains of 0.25 (IQR: 0.13 – 0.31). Approximately 80% of each strain’s gene content was shared with the other, while 18% and 24% was private to strain-6 and strain-9, respectively (Fig. 5C; Supplementary Table S3). Cross-referencing co-occurrence clusters with the estimated gene content of these strains, about 60% of clusters in each were shared, with 39% and 42% private, respectively (Fig. 5D). Of the 118 shared clusters, 12 were found in no more than two additional UCFMT strains. We hypothesize that these might indicate important physiological similarities that distinguish high-engraftment strains from the others. Among 85 genes in these shared clusters, the most common COG category annotation was X (“Mobilome”) reinforcing that phage, plasmids, and other mobile genetic elements are an important source of shared gene content across distantly related strains.

Next we sought to understand functional gene differences between the two high-engraftment strains, in particular any that might result in disparate impacts on host health. We therefore examined the unshared gene content in order to identify plausible physiological differences (Supplementary Table S3). Strikingly, strain-9 had 12 genes annotated as related to antimicrobial resistance, suggesting potential resistance to 17 different antibiotics, while strain-6 had none. Among gene co-occurrence clusters, one (labeled clust-861) is also found only in strain-9, and includes genes with homology to components of a type VI secretion system (T6SS). Most T6SSs are involved in inter-microbial competition, although a role in pathogenesis has also been described (Navarro-Garcia et al. 2019). Another cluster private to strain-9, labeled clust-37, includes genes with homology to many components of a type IV secretion system (T4SS), other secretion systems, a helicase, and a component of a toxin/anti-toxin system. Combined, these annotations suggest that the cluster may primarily reflect a mobilizable plasmid in strain-9 that is missing in strain-6. Similarly, related annotations in several clusters (clust-351, clust-352, and clust-353) have homology to genes in the *pdu*-operon. This operon encodes components of catabolic bacterial microcompartments, which are involved in various catabolic pathways, including 1,2-propanediol utilization. These co-occurrence clusters are found only in strain-9, and strain-6 is missing homology to most of the genes in the *pdu*-operon. Microcompartments and 1,2-propanediol utilization have been associated with pathogenicity in *E. coli* and other species of *Enterobacteriaceae* (Prentice 2021).

Given the presence of AMR genes and the plausible association between several co-occurrence clusters and pathogenesis, we speculate that the engraftment of *E. coli* strain-9, found in FMT samples donated by D97, could result in a less beneficial or even detrimental treatment for recipients. Similarly, the engraftment of strain-6 from D44 might contribute to the competitive exclusion of more pathogenic strains. While the previously published study found no difference in outcomes between recipients of the two donors (Smith et al. 2022b), that study may have been underpowered (n = 8 recipients for each of D44 and D97). Our computational predictions could be tested *in vitro* with isolates obtainable from archived donor materials.

## Discussion

Here we have described updates to the MIDAS v3 pangenome database and profiling software, as well as StrainPGC, a novel tool for accurate, strain-specific gene content estimation using metagenomic data. The key innovations of StrainPGC are the use of depth correlation information and selection of strain-pure samples. Together, these innovations enable StrainPGC to outperform PanPhlAn and StrainPanDA in a benchmark based on a complex, synthetic community modeled after the human gut microbiome. Combining the updated MIDAS v3 and StrainPGC in our workflow, we estimated gene content for thousands of strains in the HMP2 metagenome collection, substantially expanding on the diversity found in reference genome collections and enabling analyses of intraspecific variation without isolation or assembly. Finally, we used StrainPGC to compare the functional potential of two different strains of *E. coli* that were successfully transferred from two different donors in a clinical trial of FMT.

StrainPGC is an assembly-free method, and complements assembly-based methods—including novel, strain-aware approaches (Quince et al. 2017; Quince et al. 2021)—for gene content estimation. High-quality genome sequences enabled by laboratory isolation and culturing, as well as modern, long-read sequencing, reduce the risk of cross-mapping and remain the gold standard for comparative genomics. However, these methods are labor intensive, expensive, and often fail to capture low-abundance organisms (Chen et al. 2020). In contrast, StrainPGC offers a more accessible approach which can be applied to existing short-read metagenomic datasets. Our method identified extensive, underexplored diversity in the well-studied HMP2, demonstrating that many strains are missed by culturing and assembly-based methods. Nonetheless, both approaches are complementary: assembly-based methods contribute to the completeness and accuracy of reference databases, which in turn enhances the performance of reference-based methods like StrainPGC. Together, these diverse approaches enable comprehensive analysis of gene content variation in complex microbial communities.

Given the enormous diversity of strains found across subjects in the HMP2, the StrainPGC approach may be most useful for analyzing FMT, longitudinal, or other study designs where the same strains are expected to be found in multiple samples. While StrainPGC is specifically designed to overcome the limitations of short-read, alignment-based pangenome profiling, in particular ambiguous mapping to homologous sequences both within and across species, systematic false positive and false negative gene assignments may still occur. As a result, we caution against over-interpreting analyses that rely on directly comparing the gene content of inferred strain with reference strains. Our approach leverages strain-pure samples and compares across multiple samples with the same strain. As a result, StrainPGC will likely perform suboptimally in environments and study designs with particularly high intra-sample diversity, such as waste-water or soil microbiomes, and where fewer strain-pure samples have shared strains. This highlights an opportunity for the development of complementary tools that can handle extreme microbial diversity both within and across samples. With increasingly comprehensive pangenome reference databases, the accuracy of our approach will improve, expanding its application to other microbiomes beyond the human gut. Nonetheless, highly diverged strains may have elevated error rates due to reference database bias and it is prudent for users to ensure that their species of interest are sufficiently covered in reference sets (Zhao et al. 2023; Hovhannisyan et al. 2020). Another major barrier to interpreting gene content estimates by StrainPGC or other methods is the sparsity of robust genetic, biochemical, structural, and experimental characterization of gene products (Zhou et al. 2019). While we augmented available annotations by leveraging co-occurrence clusters to investigate epistatic and evolutionary relationships between genes—as others have done previously (Minot et al. 2021)—laboratory-based characterization is still vital.

Packaged as stand-alone software tools and integrated into an automated workflow, MIDAS v3 and StrainPGC together facilitate the broad exploration of strain-specific gene content in metagenome collections. This enables expanding surveys across additional metagenomic datasets, looking for associations between microbial strains and disease, and identifying determinants of success for FMT. Important future work also includes specializing our end-to-end workflow for environments beyond the human gut, integrating additional analyses comparing inferred strains to the reference collection, and further refining pangenome profiles based on horizontal coverage. We designed StrainPGC as part of a modular workflow that may include gene and strain information from any context. In particular, our references can be replaced with databases targeting different environments using previously released protocols (Zhao et al. 2022b; Shi et al. 2022). Thus, while we chose to focus on the human gut microbiome in this initial study, we expect that StrainPGC will be a broadly useful approach to associate genes with strains using metagenomic data from diverse environments.

## Methods

### MIDAS v3 update

Here we describe updates in MIDAS v3, including changes to the pangenome reference database construction procedure and the pangenome profiling method. Together, these updates clean, functionally annotate, and expand the phylogenetic coverage of MIDAS pangenome profiling, providing a foundation for accurately estimating and interpreting gene content across species. MIDAS v3 is available at https://github.com/czbiohub-sf/MIDAS and can be installed using conda or Docker. Compatible, pre-built MIDAS databases based on UHGG (Almeida et al. 2020) v2.0 and GTDB (Parks et al. 2021) r202 are available. We use the UHGG database throughout this work.

#### Pangenome database curation and clustering

A MIDAS v3 pangenome database can be constructed from any reference genome collection, and is composed, for each species, of predicted gene sequences from all example genomes clustered into operational gene families (OGFs) at a series of average nucleotide identity (ANI) thresholds. For clarity, we have referred to these OGFs simply as genes in the main text. In order to minimize the impacts of inter-and intra-specific cross-mapping on pangenome profiling, which can be major problems for gene databases constructed with MAGs, we made major changes to the clustering and curation pipeline. In this MIDAS update, described below, we sought to minimize the impact of fragmented gene sequences, spurious gene calls, chimeric assemblies, and redundant OGFs resulting from these errors (Li et al. 2014; Hyatt et al. 2012; Dimonaco et al. 2021).

For each species, for each reference genome in the source genome collection, genes were predicted by Prokka v1.14.6 (Seemann 2014), wrapping Prodigal v2.6.3 (Hyatt et al. 2010). Gene sequences less than 200 bp or with ambiguous bases (anything but A, C, G, or T) were removed. Then, the remaining sequences were dereplicated by clustering at a 99% ANI threshold using VSEARCH v2.23.0 (Rognes et al. 2016), with the longest sequence initially assigned as the representative sequence for the cluster. Next, in order to identify and remove additional cases of fragmented genes, we applied CD-HIT v4.8.1 (Fu et al. 2012) (using options `-c 1 -aS 0.9 -G 0 -g 1 -AS 180`); when a shorter representative sequence had perfect identity over ≥ 90% of length to a longer sequence, the two clusters were merged, and the longer sequence was assigned as representative. Short gene sequences predicted on the opposite strand, a known complication (Trimble et al. 2012), were also merged in this way.

Having dereplicated and cleaned gene sequences, we further clustered representative sequences into OGFs using VSEARCH, defining final OGF clusters at thresholds between 95% and 75% ANI.

#### Pangenome database annotation

Next, we annotated sequences using a variety of tools. We ran EggNOG mapper v2.1.12 (Cantalapiedra et al. 2021) on dereplicated genes to identify homology relative to several commonly used gene orthologies: COGs, EggNOG OGs, and KOs. ResFinder v4.4.2 (Florensa et al. 2022), geNomad v1.7.4 (Camargo et al. 2023) and MobileElementFinder v1.1.2 (Johansson et al. 2020) were run directly on contigs of each reference genome to identify AMR, phage, plasmid, and mobile element associated regions, and these annotations were transferred onto predicted genes based on overlapping coordinates.

While annotations are performed on genomic sequences or dereplicated gene sequences, interpretation of estimated gene content requires annotations at the OGF level. We therefore implemented a voting procedure intended to enable the transfer of annotations from gene sequences to gene clusters. For OGFs at each ANI level, we calculated the fraction of genes in each cluster annotated as an AMR gene, phage-associated, plasmid-associated, or mobile element-associated. In this way, users can identify annotations robustly associated with genes of interest.

#### Alignment and gene depth estimation

For pangenome profiling, the MIDASDB representative gene sequences from selected species (i.e. dereplicated at the 99% ANI level) are compiled into an index for alignment and quantification. At this stage, we apply additional filtering to the set of representative sequences, which we refer to as “pruning”, with the goal of speeding up alignment and improving quantification by reducing the rate of cross-mapping within and between species. First, we remove representative sequences that are less than 50% of the median length in the 95% ANI cluster, as these are more likely to be truncated genes resulting from assembly fragmentation. Second, for species with more than 10 reference genomes, we remove representative sequences where their 75% ANI clusters had only one member, as these are more likely to be spurious gene calls or contamination resulting from chimeric assembly. Finally, an alignment index is constructed from the remaining representative sequences, and reads are mapped using Bowtie2 (Langmead and Salzberg 2012).

Pangenome profiling with MIDAS v3 proceeds through four stages: (1) building a reference index as described above, (2) alignment of reads to the reference index, (3) calculation of the mean depth across the length of the representative sequence, and then (4) summation of representative sequence depths into clusters in order to estimate the total depth of the OGF at the chosen ANI threshold.

### Shotgun metagenomes

All shotgun metagenomes analyzed in this work are publicly available as SRA BioProjects, including the HMP2 (PRJNA398089), UCFMT (PRJNA737472), and the hCom2 samples used for benchmarking (PRJNA885585). The HMP2 metagenomes already had human reads removed and quality control procedures previously applied. UCFMT metagenomes were filtered for human reads, deduplicated, adapter trimmed, and quality trimmed, as previously described in (Smith et al. 2022b). The hCom2 metagenomes were processed in the same way, except that human read removal was skipped because the data was collected *in vitro*.

### Integrated analysis workflow

#### Pangenome profiling

For the work presented here, we ran MIDAS v3 Using Bowtie2 v2.5.1 throughout, a single reference index was built for 627 species using `midas build_bowtie2db --prune_centroids --remove_singleton`. Paired-end reads for each sample were aligned to this index using `midas run_genes --aln_speed sensitive --aln_extra_flags ’--mm --ignore-quals’ --total_depth 0`. To maximize our sensitivity to divergent strains and at low abundance we did not use any of MIDAS’s default filters in calculating depths. Instead, mean mapping depth was calculated using `samtools depth` and summed up at the 75% ANI OGF level.

#### Reference genomes and species marker genes

High-quality reference genomes in the UHGG were defined as those with estimated completeness of > 90% and contamination of < 5%. OGFs found in > 95% of high-quality reference genomes were selected as species marker genes and were used for species depth estimation, quality control, and downstream analyses.

A list of the marker genes used for each of the 627 species analyzed in this study are distributed with the StrainPGC software.

#### SNP profiling

SNP profiles were obtained from metagenomes using GT-Pro v1.0.1 (Shi et al. 2022) and the default database, which was built using UHGG v1.0. GT-Pro was run on preprocessed reads, and counts from forward and reverse reads were summed. The resulting SNP profile matrix, a three-dimensional array of counts indexed by sample, genotyped position, and allele (reference or alternative), is the core input for StrainFacts (Smith et al. 2022a).

An analogous approach was used to obtain SNP genotypes for genomic sequence. Specifically, for both reference and benchmarking genomes, contigs were fragmented into 500 bp tiles with 31 bp of overlap and used as input to GT-Pro. We filtered out tallies for SNP sites that did not match the expected species.

#### Strain tracking and genotyping

For each species, SNP profiles obtained from GT-Pro were filtered to remove low-depth samples (those with < 5% of positions observed). For the HMP2 and UCFMT datasets, low polymorphism positions (minority allele observed in < 5% of samples) were also removed. However, this latter filter was not applied to the synthetic community since many species had only one strain. Strain genotypes and proportions were estimated with StrainFacts v0.6.0, using the updated Model 4 and a number of strains set as *n*^0.85^ where *n* is the number of samples. For the vast majority of species, this model was fit using a single, standardized set of hyperparameters: --optimizer-learning-rate 0.05 --min-optimizer-learning-rate 1e-2 --hyperparameters gamma_hyper=1e-15 pi_hyper=1e-2 pi_hyper2=1e-2 rho_hyper=1.0 rho_hyper2=1.0 --anneal-hyperparameters gamma_hyper=0.999 --anneal-steps 120000. However, for seven species (species IDs: sp-100076, sp-101302, sp-101306, sp-101704, sp-102478, sp-103456, sp-103683), amended hyperparameters were found to perform better: `gamma_hyper=1e-10 pi_hyper=1e-3 pi_hyper2=1e-3 gamma_hyper=1e-1 rho_hyper=10.0 rho_hyper2=10.0 -- anneal-steps 20000`.

Each strain-pure set was defined as those samples where StrainFacts estimated it to be > 95% of the species. For analyses requiring estimated genotypes, we used a consensus genotype for each strain, pooling all samples in the strain-pure set. Based on this pooling, the consensus genotype for each strain was the majority allele at each position. Positions with unexpectedly high counts of the minor allele (≥ 10%)—which suggests issues with genotyping—were masked. Similarly, positions without any observed alleles were also masked in subsequent comparisons. Likewise, SNPs in reference and benchmark genotypes where neither allele was observed were masked in downstream analyses. We selected this as a more conservative approach compared to directly using the genotypes estimated by StrainFacts. All pairwise dissimilarities between inferred strain, reference, and benchmark genotypes were calculated as the masked Hamming distance, with a pseudocount of 1 added, i.e.: 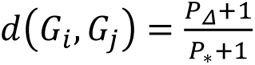 where *P_Δ_* is the number of positions with different allele and *P_*_* is the number of unmasked positions.

Note that this measure of genetic distance is related to but not equivalent to the complement of the core genome average nucleotide identity (“ANI dissimilarity”: 1 − ANI), since it is based on only known polymorphic sites in the core genome, and the actual ANI dissimilarity—which includes many non-polymorphic sites in the denominator, as well—is likely to be much smaller.

#### StrainPGC

For each species we estimated gene content across strains with StrainPGC v0.1.0, providing the three required inputs: (1) the list of species marker gene IDs from the MIDASDB, (2) the strain pure sets derived from StrainFacts, and (3) pangenome profiles from MIDAS as the three inputs.

StrainPGC estimates the depth of each species in each sample as the 15%-trimmed mean depth across all species marker genes, i.e., the mean depth of species marker genes excluding those genes with the 15% highest and lowest depth. Species-free samples were defined as those with an estimated species depth of < 0.0001x. Genes were selected using a depth ratio threshold of 0.2 and a correlation threshold of 0.4 in order to strike a balance between sensitivity and specificity, while slightly favoring false negatives over false positives (see Supplementary Figure S3).

#### Gene family annotation

To facilitate functional interpretation, we extended the voting procedure used for the MIDASDB to EggNOG mapper annotations, which include COGs, COG categories, EggNOG OGs, KOs, and KEGG Modules. We augmented the COG categories assigned by EggNOG mapper with additional categories available from https://ftp.ncbi.nlm.nih.gov/pub/COG/COG2020/data. Since annotations were performed on representative sequences for each dereplicated gene (99% ANI cluster), we first transferred specific annotations to all cluster members. Annotations within each gene (75% ANI cluster) were then counted as votes. Any annotations possessed by > 50% of member sequences were assigned to the gene family as a whole. Note that while the annotation voting for the MIDASDB, described above, operates on binary annotations (e.g., it is or is not a phage gene), this additional voting procedure was performed for individual annotations (e.g., a specific COG or AMR reference accession).

### Downstream analysis

#### Benchmarking gene content estimation performance

We benchmarked the performance of gene content estimates from StrainPGC, PanPhlAn, and StrainPanDA, using publicly available, high-quality strain genomes and metagenomes from experimental treatments of the hCOM2 synthetic community (Jin et al. 2023). From the 117 inoculated strains, we excluded genomes from evaluation if: (1) when running GT-Pro directly on their genome sequence, less than 50% of identified SNPs were from the same species, or (2) the species had no depth across metagenomes, as estimated from mean marker gene depth.

In the remaining 97 strain genomes we identified gene sequences with Prodigal v2.6.3 (Hyatt et al. 2012) (masking ambiguous bases and using the `meta` procedure), translated them with codon table 11, and annotated them with EggNOG mapper version 2.1.10. The ground-truth annotations used to assess performance were defined as the complete set of all EggNOG OGs assigned to all genes in the ground-truth genome. These were compared to the complete set of OG annotations in each inferred strain’s estimated gene content.

In order to select which inferred strain to compare to each benchmark genome, the GT-Pro genotype of the ground-truth genome was compared to all strain-pure sample consensus genotypes, and the best match was identified based on the smallest masked hamming distance. For each benchmark genome, we calculated the precision, recall, and F1 score for this best match.

Both PanPhlAn and StrainPanDA are packaged with their own pangenome databases and profiling scripts. However, in order to compare the core algorithms directly, the same MIDAS pangenome profiles were provided as input to all three tools. Both alternative tools have several parameters that control when they fail to run on low sequencing depth datasets. Since, for some species, the use of default parameter values results in a runtime exception, we adjusted these parameters to be much more lenient. For PanPhlAn, we used the flags: `--left_max 1000000 --right_min 0 --min_coverage 0`. For StrainPanDA, we made modifications to the code (see https://github.com/bsmith89/StrainPanDA) and used the flags `--mincov 10 –minfrac 0.9 --minreads 1e6 --minsamples 1`. We also fixed the number of latent strains to 6 using `--max_rank 6 --rank 6` for all runs. For PanPhlAn and StrainPanDA, the inferred strain with the highest F1 score was used for performance comparisons.

#### Inferred strain quality filtering

For analysis of the HMP2 and UCFMT datasets—but not performance benchmarking—strains were filtered to remove those likely to be low accuracy. Strains with fewer than 100 unmasked positions in their consensus genotype were included in benchmarking but excluded from all other analyses. This criterion *a priori* excludes 19 of the 627 species profiled in this work. For analyses of gene content, strains with an estimated depth of < 1x across all strain-pure samples were also excluded. Finally, strains with < 95% of species genes or with a standard deviation in the log10-transformed depth ratio across selected genes of > 0.25 were flagged as low quality and removed.

#### Analysis of species and strain diversity

The species phylogeny in Fig. 2A and Fig. 3E was obtained directly from the UHGG https://ftp.ebi.ac.uk/pub/databases/metagenomics/mgnify_genomes/human-gut/v2.0.2/phylogenies/bac120_iqtree.nwk.

For the analysis of strain distribution in the HMP2, strain depth was estimated as the product of the estimated species depth and estimated strain fraction. All strains with depth > 0.1x were considered to be “present” in a sample. The number of strains in each subject was calculated as the total number of strains present in any of that subject’s samples. For shared-strain analysis (Fig. 3C), samples with fewer than 10 strains present of any species were excluded from analysis, as this removed several samples with anomalously low diversity.

Gene content was compared using the cosine dissimilarity. For comparisons between inferred strains and references, the inferred strains’ gene content was first batch corrected by subtracting the difference in means (i.e., the difference in prevalence).

#### Pangenome Analyses

To calculate the correlation between gene prevalence in reference genomes and inferred strains we first removed genes that were very rare (< 1%) in both.

Genes found in no more than one or missing from no more than one genome were excluded from clustering analysis. The remaining genes were then hierarchically clustered based on their correlation across inferred strains using the average-neighbor method at a correlation threshold of 0.9. Only clusters with more than one member were kept.

To analyze the clumping of related genes in co-occurrence clusters, we considered annotations of (1) individual KEGG modules and (2) binary classification of genes as phage and/or plasmid. For each co-occurrence cluster, we took the maximum count for any one annotation. To estimate a distribution under the null, we permuted cluster labels within species before again collecting the maximum counts across clusters. Significance was tested by comparing the number of clusters with ≥3 related annotations to the null.

For analysis of the UCFMT *E. coli* strains, shell genes and co-occurrence clusters were defined using the HMP2 inferred strains, not *de novo*.

## Software availability

StrainPGC is freely available at https://github.com/bsmith89/StrainPGC. Code and metadata needed to replicate our analyses and plots are available at https://github.com/bsmith89/StrainPGC-manuscript.

## Acknowledgments

This work was funded by a NHLBI grant #HL160862, the Chan Zuckerberg Biohub San Francisco, Gladstone Institutes, and the Sam Simeon Fund. BJS was supported by a Computational Innovation Postdoctoral Fellowship from the Noyce Initiative for Digital Transformation in Computational Biology and Health Data Science. JA was supported by funding from the Kenneth Rainin Foundation and the Crohn’s and Colitis Foundation. The authors thank Françoise Chanut for extensive editorial support.

## Author Contributions

BJS: Conceptualization, Methodology, Software, Formal Analysis, Investigation, Writing – Original Draft, Writing – Review & Editing, Visualization; CZ: Conceptualization, Methodology, Software, Writing – Original Draft, Writing – Review & Editing; VD: Writing – Review & Editing, Visualization; XJ: Writing – Original Draft, Writing – Review & Editing, Visualization; LZ: Writing – Review & Editing; SS: Software, Validation, Writing – Review & Editing; JA: Data Curation, Writing – Review & Editing; ES: Supervision, Funding Acquisition, Writing – Review & Editing; KP: Conceptualization, Methodology, Investigation, Resources, Writing – Original Draft, Writing – Review & Editing, Supervision, Funding Acquisition.

## Supplementary Materials

### Extended hCom2 benchmarking results

**Figure S1:**
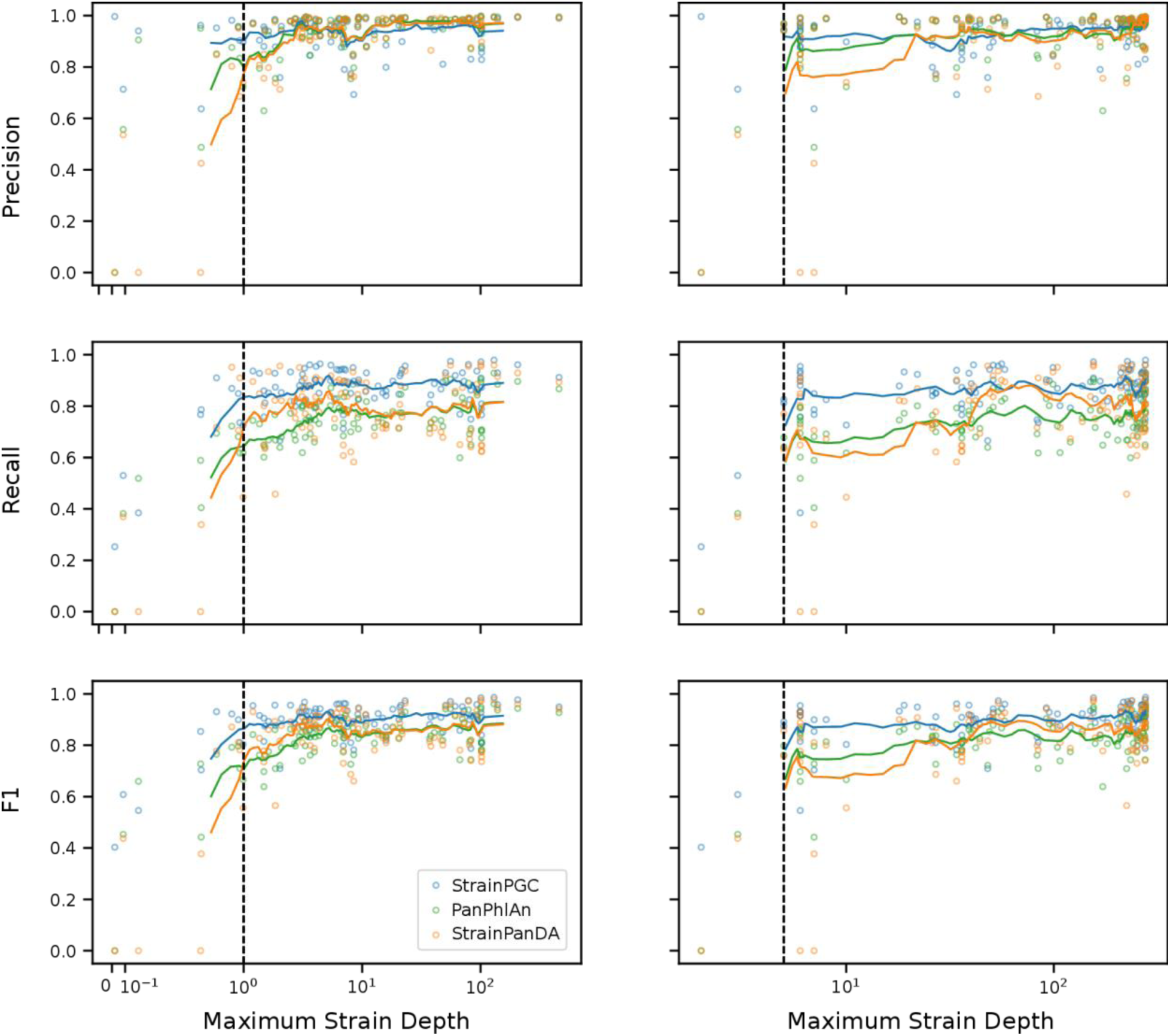
Relationship between sequencing depth or number of samples and the accuracy of gene content estimation. Points represent the performance of each tool (colors) on each of the 97 benchmark strains. For the left column, the x-axis is the maximum estimated depth of the genotype-matched strain across strain-pure samples, and for the right column it is the total number of strain-pure samples identified for that strain. Trend lines are a rolling average over the 10 nearest points. The dotted vertical line indicates the 1x depth and 5 strain-pure samples, after which the mean performance stabilizes for StrainPGC.

### Extended pangenome results

**Figure S2:**
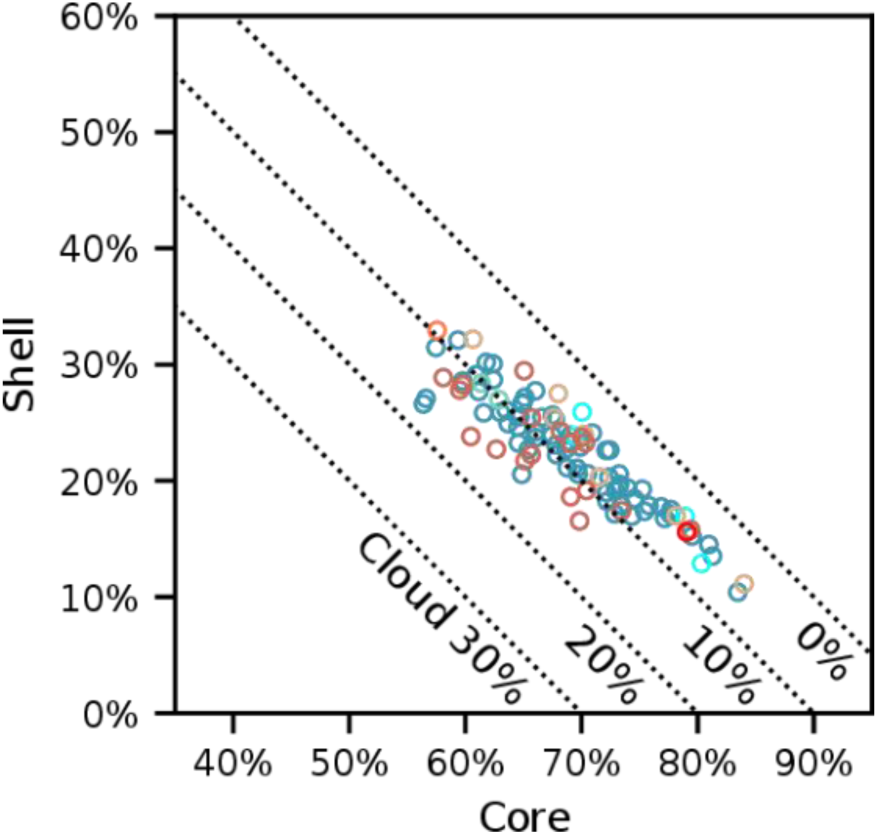
Per-genome core, shell, and cloud gene fractions in reference genomes. Equivalent results to Fig. 4B, here calculated using reference genomes for comparison to StrainPGC-based gene content estimates.

### Simulated *E. coli* spike-in validation

We performed an additional benchmarking study to validate our approach in datasets with substantially more strain-diversity, for strains with more divergence from the reference set, and with a limited number of strain-pure samples. To keep the simulated metagenomeic data as realistic as possible, we opted to construct samples with novel strains by “spiking” simulated reads from recently sequenced isolates into real metagenomes from the HMP2 study. Due to an abundance of studies with wild *E. coli* isolates, and our particular focus on this species throughout, we identified five novel *E. coli* genomes from a recently published project (Davidova-Gerzova et al. 2023). These isolates varied greatly in their relatedness to the UHGG reference genomes, including very distantly related strains with a genotype dissimilarity of 0.077. These strains are as novel relative to the reference database as would be expected for *E. coli* found in the human gut; only 0.8% of UHGG genomes had a closest match genotype-dissimilarity of more than 0.077.

We selected five HMP2 samples, all from one subject (C3022), where *E. coli* was not detected. Into these, we spiked-in simulated reads at 1x, 2x, 4x, 8x, and 16x depths with a separate set of reads for each strain. We combined all 25 of these additional, synthetic samples with the full HMP2 dataset, and then re-ran our integrated workflow. We matched the inferred strains to each of the ground-truth genomes based on genotype similarity and evaluated the StrainPGC gene content estimates as in the hCom2 benchmark.

We found that the performance of StrainPGC in these simulations with non-reference *E. coli* genomes is consistent with the overall performance on the hCom2 (synthetic community) benchmark. This is despite the fact that the metagenomes were much more complex and some strains were more dissimilar to the closest reference genome. Specifically, we found a median F1 score across all strains of 0.92, equivalent to the median F1 of 0.91 from the hCom2 benchmark. Interestingly, we do not find a negative relationship between the divergence of the benchmark genome and performance. StrainPGC performance was nearly equivalent for the least diverged (F1 of 0.89) and most diverged genomes (F1 of 0.92). We conclude that it is reasonable to expect similar performance for other strains and datasets, even when the number of strains for a species is large and when strains are more diverged from the reference database.

**Supplementary Table S2:**
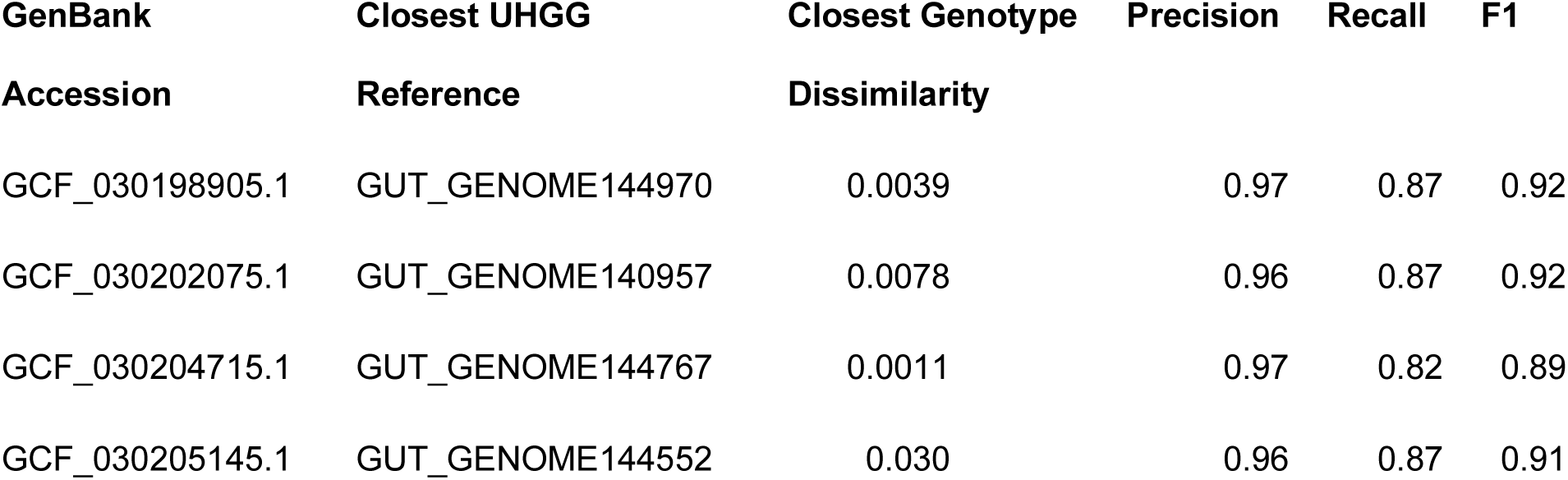
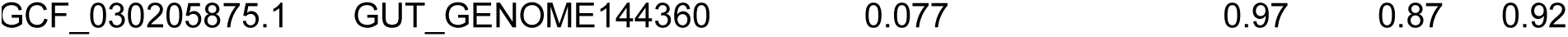
Performance on five E. coli genomes in an in silico spike-in experiment.

### Sensitivity of StrainPGC performance to depth ratio and correlation score thresholds

**Figure S3:**
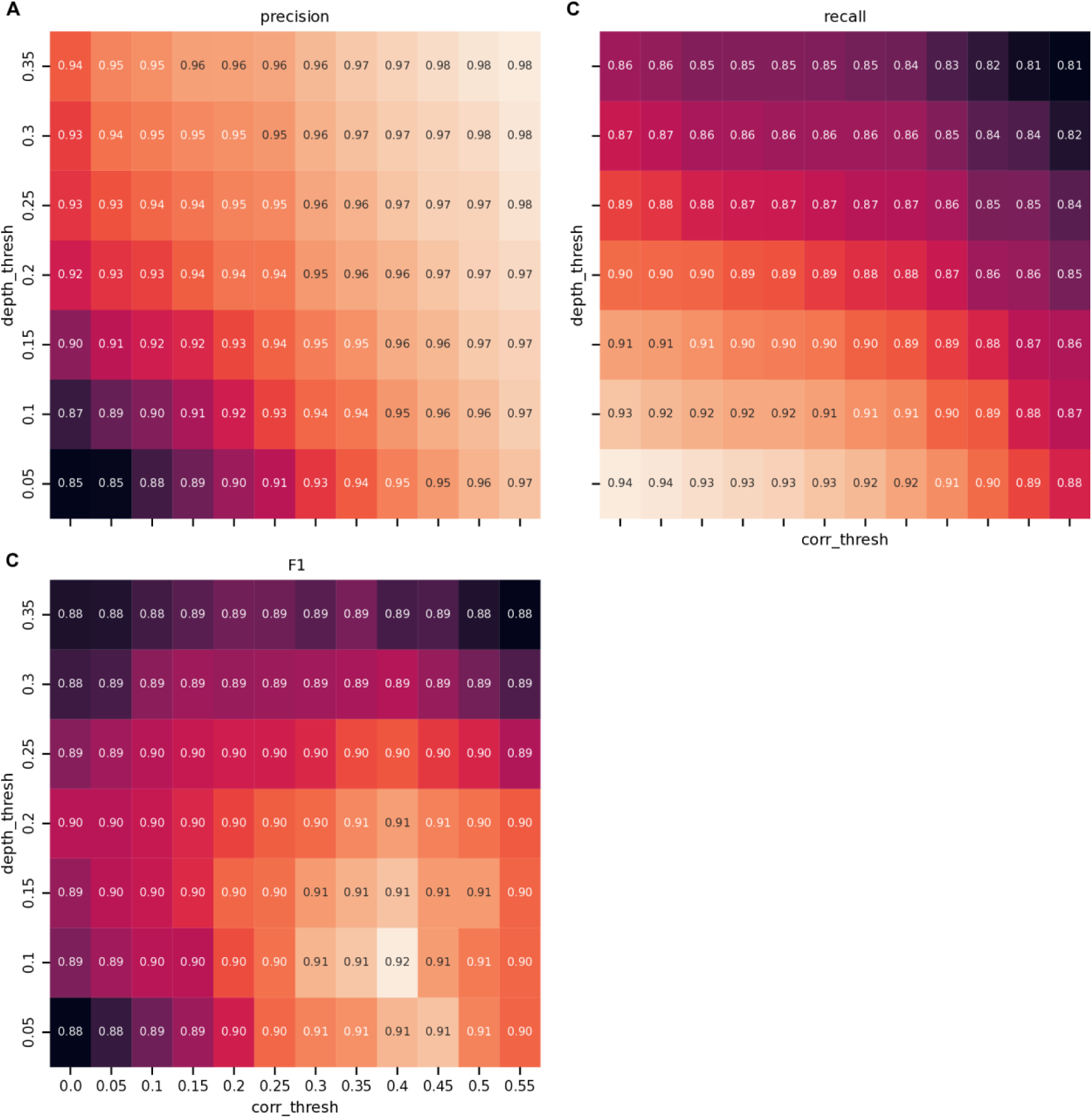
Threshold depth ratio and correlation score parameter search. Median performance across 97 hCom2 benchmark genomes at every combination of 11 correlation score thresholds (x-axis) and 7 depth ratio thresholds (y-axis). Panels represent median precision (A), recall (B), and F1 score (C). The best performance (F1 score) was achieved at a depth ratio threshold of 0.1 and correlation threshold of 0.40. We used a slightly more conservative depth threshold of 0.2 throughout the rest of this work, which decreased the median F1 score negligably from 0.916 to 0.908.

## References

1. Almeida A, Nayfach S, Boland M, Strozzi F, Beracochea M, Shi ZJ, Pollard KS, Sakharova E, Parks DH, Hugenholtz P, et al. 2020. A unified catalog of 204,938 reference genomes from the human gut microbiome. Nat Biotechnol 39: 105–114.

2. Beghini F, McIver LJ, Blanco-Míguez A, Dubois L, Asnicar F, Maharjan S, Mailyan A, Manghi P, Scholz M, Thomas AM, et al. 2021. Integrating taxonomic, functional, and strain-level profiling of diverse microbial communities with bioBakery 3. eLife 10: e65088.

3. Blanco-Míguez A, Beghini F, Cumbo F, McIver LJ, Thompson KN, Zolfo M, Manghi P, Dubois L, Huang KD, Thomas AM, et al. 2023. Extending and improving metagenomic taxonomic profiling with uncharacterized species using MetaPhlAn 4. Nat Biotechnol 41: 1633–1644.

4. Blount ZD. 2015. The unexhausted potential of E. coli. eLife 4: e05826.

5. Camargo AP, Roux S, Schulz F, Babinski M, Xu Y, Hu B, Chain PSG, Nayfach S, Kyrpides NC. 2023. Identification of mobile genetic elements with geNomad. Nat Biotechnol 1–10.

6. Cantalapiedra CP, Hernández-Plaza A, Letunic I, Bork P, Huerta-Cepas J. 2021. eggNOG-mapper v2: Functional Annotation, Orthology Assignments, and Domain Prediction at the Metagenomic Scale. Molecular Biology and Evolution 38: 5825–5829.

7. Carr R, Shen-Orr SS, Borenstein E. 2013. Reconstructing the Genomic Content of Microbiome Taxa through Shotgun Metagenomic Deconvolution. PLoS Comput Biol 9: e1003292.

8. Carrow HC, Batachari LE, Chu H. 2020. Strain diversity in the microbiome: Lessons from Bacteroides fragilis. PLoS Pathog 16: e1009056.

9. Chen L-X, Anantharaman K, Shaiber A, Eren AM, Banfield JF. 2020. Accurate and complete genomes from metagenomes. Genome Res 30: 315–333.

10. Cheng AG, Ho P-Y, Aranda-Díaz A, Jain S, Yu FB, Meng X, Wang M, Iakiviak M, Nagashima K, Zhao A, et al. 2022. Design, construction, and in vivo augmentation of a complex gut microbiome. Cell 185: 3617–3636.e19.

11. Davidova-Gerzova L, Lausova J, Sukkar I, Nesporova K, Nechutna L, Vlkova K, Chudejova K, Krutova M, Palkovicova J, Kaspar J, et al. 2023. Hospital and community wastewater as a source of multidrug-resistant ESBL-producing Escherichia coli. Front Cell Infect Microbiol 13.

12. Dimonaco NJ, Aubrey W, Kenobi K, Clare A, Creevey CJ. 2021. No one tool to rule them all: Prokaryotic gene prediction tool annotations are highly dependent on the organism of study. Bioinformatics 38: 1198–1207.

13. Florensa AF, Kaas RS, Clausen PTLC, Aytan-Aktug D, Aarestrup FM. 2022. ResFinder – an open online resource for identification of antimicrobial resistance genes in next-generation sequencing data and prediction of phenotypes from genotypes. Microb Genom 8: 000748.

14. Fu L, Niu B, Zhu Z, Wu S, Li W. 2012. CD-HIT: Accelerated for clustering the next-generation sequencing data. Bioinformatics 28: 3150–3152.

15. Hovhannisyan H, Hafez A, Llorens C, Gabaldón T. 2020. CROSSMAPPER: Estimating cross-mapping rates and optimizing experimental design in multi-species sequencing studies. Bioinformatics 36: 925–927.

16. Hu H, Tan Y, Li C, Chen J, Kou Y, Xu ZZ, Liu Y, Tan Y, Dai L. 2022. StrainPanDA: Linked reconstruction of strain composition and gene content profiles via pangenome-based decomposition of metagenomic data. iMeta 1: e41.

17. Hyatt D, Chen G-L, LoCascio PF, Land ML, Larimer FW, Hauser LJ. 2010. Prodigal: Prokaryotic gene recognition and translation initiation site identification. BMC Bioinformatics 11: 119.

18. Hyatt D, LoCascio PF, Hauser LJ, Uberbacher EC. 2012. Gene and translation initiation site prediction in metagenomic sequences. Bioinformatics 28: 2223–2230.

19. Jin X, Yu FB, Yan J, Weakley AM, Dubinkina V, Meng X, Pollard KS. 2023. Culturing of a complex gut microbial community in mucin-hydrogel carriers reveals strain-and gene-associated spatial organization. Nat Commun 14: 3510.

20. Joglekar P, Sonnenburg ED, Higginbottom SK, Earle KA, Morland C, Shapiro-Ward S, Bolam DN, Sonnenburg JL. 2018. Genetic Variation of the SusC/SusD Homologs from a Polysaccharide Utilization Locus Underlies Divergent Fructan Specificities and Functional Adaptation in Bacteroides Thetaiotaomicron Strains. mSphere 3.

21. Johansson MHK, Bortolaia V, Tansirichaiya S, Aarestrup FM, Roberts AP, Petersen TN. 2020. Detection of mobile genetic elements associated with antibiotic resistance in Salmonella Enterica using a newly developed web tool: MobileElementFinder. J Antimicrob Chemother 76: 101–109.

22. Langmead B, Salzberg SL. 2012. Fast gapped-read alignment with Bowtie 2. Nat Methods 9: 357–359.

23. Li J, Jia H, Cai X, Zhong H, Feng Q, Sunagawa S, Arumugam M, Kultima JR, Prifti E, Nielsen T, et al. 2014. An integrated catalog of reference genes in the human gut microbiome. Nat Biotechnol 32: 834–841.

24. Lloyd-Price J, Mahurkar A, Rahnavard G, Crabtree J, Orvis J, Hall AB, Brady A, Creasy HH, McCracken C, Giglio MG, et al. 2017. Strains, functions and dynamics in the expanded Human Microbiome Project. Nature 550: 61–66.

25. Milanese A, Mende DR, Paoli L, Salazar G, Ruscheweyh H-J, Cuenca M, Hingamp P, Alves R, Costea PI, Coelho LP, et al. 2019. Microbial abundance, activity and population genomic profiling with mOTUs2. Nat Commun 10: 1014.

26. Minot SS, Barry KC, Kasman C, Golob JL, Willis AD. 2021. Geneshot: Gene-level metagenomics identifies genome islands associated with immunotherapy response. Genome Biol 22: 135.

27. Mölder F, Jablonski KP, Letcher B, Hall MB, Tomkins-Tinch CH, Sochat V, Forster J, Lee S, Twardziok SO, Kanitz A, et al. 2021. Sustainable data analysis with Snakemake.

28. Navarro-Garcia F, Ruiz-Perez F, Cataldi Á, Larzábal M. 2019. Type VI Secretion System in Pathogenic Escherichia coli: Structure, Role in Virulence, and Acquisition. Front Microbiol 10: 1965.

29. Nayfach S, Rodriguez-Mueller B, Garud N, Pollard KS. 2016. An integrated metagenomics pipeline for strain profiling reveals novel patterns of bacterial transmission and biogeography. Genome Res 26: 1612–1625.

30. Pakbin B, Brück WM, Rossen JWA. 2021. Virulence Factors of Enteric Pathogenic Escherichia coli: A Review. IJMS 22: 9922.

31. Parks DH, Chuvochina M, Rinke C, Mussig AJ, Chaumeil P-A, Hugenholtz P. 2021. GTDB: An ongoing census of bacterial and archaeal diversity through a phylogenetically consistent, rank normalized and complete genome-based taxonomy. Nucleic Acids Research 50: D785–D794.

32. Plaza Oñate F, Le Chatelier E, Almeida M, Cervino ACL, Gauthier F, Magoulès F, Ehrlich SD, Pichaud M. 2018. MSPminer: Abundance-based reconstitution of microbial pan-genomes from shotgun metagenomic data. Bioinformatics 35: 1544–1552.

33. Prentice MB. 2021. Bacterial microcompartments and their role in pathogenicity. Current Opinion in Microbiology 63: 19–28.

34. Proctor LM, Creasy HH, Fettweis JM, Lloyd-Price J, Mahurkar A, Zhou W, Buck GA, Snyder MP, Strauss JF, Weinstock GM, et al. 2019. The Integrative Human Microbiome Project. Nature 569: 641–648.

35. Quince C, Delmont TO, Raguideau S, Alneberg J, Darling AE, Collins G, Eren AM. 2017. DESMAN: A new tool for de novo extraction of strains from metagenomes. Genome Biol 18: 1– 22.

36. Quince C, Nurk S, Raguideau S, James R, Soyer OS, Summers JK, Limasset A, Eren AM, Chikhi R, Darling AE. 2021. STRONG: Metagenomics strain resolution on assembly graphs. Genome Biology 22: 214.

37. Ray S, Das S, Suar M. 2017. Molecular Mechanism of Drug Resistance. In Drug Resistance in Bacteria, Fungi, Malaria, and Cancer (eds. G. Arora, A. Sajid, and V.C. Kalia), pp. 47–110, Springer International Publishing, Cham.

38. Rognes T, Flouri T, Nichols B, Quince C, Mahé F. 2016. VSEARCH: A versatile open source tool for metagenomics. PeerJ 4: e2584.

39. Seemann T. 2014. Prokka: Rapid prokaryotic genome annotation. Bioinformatics 30: 2068– 2069.

40. Shi ZJ, Dimitrov B, Zhao C, Nayfach S, Pollard KS. 2022. Fast and accurate metagenotyping of the human gut microbiome with GT-Pro. Nat Biotechnol 40: 507–516.

41. Smith BJ, Li X, Shi ZJ, Abate A, Pollard KS. 2022a. Scalable Microbial Strain Inference in Metagenomic Data Using StrainFacts. Front Bioinform 2.

42. Smith BJ, Piceno Y, Zydek M, Zhang B, Syriani LA, Terdiman JP, Kassam Z, Ma A, Lynch SV, Pollard KS, et al. 2022b. Strain-resolved analysis in a randomized trial of antibiotic pretreatment and maintenance dose delivery mode with fecal microbiota transplant for ulcerative colitis. Sci Rep 12: 5517.

43. Trimble WL, Keegan KP, D’Souza M, Wilke A, Wilkening J, Gilbert J, Meyer F. 2012. Short-read reading-frame predictors are not created equal: Sequence error causes loss of signal. BMC Bioinformatics 13: 183.

44. Yang C, Mogno I, Contijoch EJ, Borgerding JN, Aggarwala V, Li Z, Siu S, Grasset EK, Helmus DS, Dubinsky MC, et al. 2020. Fecal IgA Levels Are Determined by Strain-Level Differences in Bacteroides ovatus and Are Modifiable by Gut Microbiota Manipulation. Cell Host & Microbe 27: 467–475.e6.

45. Zhao C, Dimitrov B, Goldman M, Nayfach S, Pollard KS. 2022a. MIDAS2: Metagenomic Intra-species Diversity Analysis System. Bioinformatics 39: btac713.

46. Zhao C, Goldman M, Smith BJ, Pollard KS. 2022b. Genotyping Microbial Communities with MIDAS2: From Metagenomic Reads to Allele Tables. Current Protocols 2: e604.

47. Zhao C, Shi ZJ, Pollard KS. 2023. Pitfalls of genotyping microbial communities with rapidly growing genome collections. Cell Systems 14: 160–176.e3.

48. Zhou N, Jiang Y, Bergquist TR, Lee AJ, Kacsoh BZ, Crocker AW, Lewis KA, Georghiou G, Nguyen HN, Hamid MN, et al. 2019. The CAFA challenge reports improved protein function prediction and new functional annotations for hundreds of genes through experimental screens. Genome Biol 20: 244.

